# A natural cyclic peptide valinomycin enhances plant innate immunity

**DOI:** 10.1101/2023.10.25.563898

**Authors:** Nayeon Yoo, Ji Eun Kang, Da-Ran Kim, Huiwon Lee, Dohee Ko, Youn-Sig Kwak, Eui-Hwan Chung

## Abstract

Various natural compounds as alternative agents for the chemical management of plant diseases have long been proposed. Valinomycin, a *Streptomyces*-derived cyclic peptide, acts as an antifungal agent against several plant pathogenic fungi, including *Botrytis cinerea*. Here, we report the novel function of valinomycin, plant immune-boosting. Valinomycin potentiates pattern-triggered immunity (PTI) and effector-triggered immunity (ETI) in *Arabidopsis*, leading to enhanced resistance against bacterial speck disease locally and systemically. Moreover, this plant immune-boosting activity of valinomycin is associated with plant hormonal signaling. Thus, we propose that valinomycin harbors potential as a biocontrol agent suppressing complex pathogen infections, such as bacteria and fungi.

## INTRODUCTION

Plants are sessile organisms incapable of escaping complex pathogen attacks. Unlike animals, plants rely solely on two-tiered innate immunity composed of pattern-triggered immunity (PTI) and effector-triggered immunity (ETI) (Jones & Dangl, 2006). During PTI, pattern-recognition receptors (PRRs) recognize pathogen-associated molecular patterns (PAMPs). Pathogens secrete effectors to dampen PTI response in plants. These effectors can be perceived by intracellular nucleotide-binding leucine-rich repeat receptors (NLRs) directly and indirectly, leading to the activation of ETI (Dodds & Rathjen, 2010; Jones & Dangl, 2006). PTI and ETI share common immune outputs, such as reactive oxygen species (ROS) burst and mitogen-activated protein kinase (MAPK) activation (Chang et al., 2022). Intriguingly, recent studies reported mutual potentiation of PTI and ETI for more effective and heightened plant immunity (Pruitt et al., 2021).

Plant disease management has relied on chemicals, primarily antibiotic agents that directly eradicate pests. However, antibiotic resistance emerged as a major concern as pesticides gradually lost their efficacy. Thus, new research directions focusing on enhancing plant immunity and suppressing pathogen virulence have been proposed as promising alternatives (Dickey et al., 2017; Kang et al., 2022). With these aims, natural compounds inducing plant resistance have been widely identified, including β-aminobutyric acid, resveratrol oligomers, and bacterial PAMPs (Cohen et al., 2016; Kang et al., 2022; Zipfel et al., 2004). Moreover, various plant-derived natural compounds, including hopeaphenol, were identified as anti-virulence agents against pathogens (Kang, Hwang, et al., 2022; Silva et al., 2016).

Here, we explored the impact of *Streptomyces*-producing valinomycin on plant immunity by conducting a comprehensive examination of PTI and ETI outcomes following valinomycin treatment. Our findings revealed that valinomycin modulates mutual potentiation of PTI and ETI locally, possibly resulting in improved systemic resistance. Thus, this study offers a new perspective on characterizing natural immune-boosting compounds, along with insights into the underlying mechanism.

## RESULTS AND DISCUSSION

### Valinomycin facilitates more robust PTI and ETI responses

Valinomycin, isolated from *Streptomyces bacillaris* sp. S8 culture extract (Fig, S1A - E), is an ionophore that leads to a collapse of the proton motive force and disrupts bacterial cell membranes, while its function against plant bacterial pathogens was not yet fully investigated (Cooper et al., 2012; Farha et al., 2013). We first determined a reasonable valinomycin concentration for further experiments by minimal inhibitory concentration (MIC) test (Fig. S1F). Valinomycin completely inhibited fungal growth of *Botrytis cinerea* (*B. cinerea*) at the concentration above the MIC value (≥ 10 μM), consistent with previous finding (Park et al., 2008). Valinomycin exhibited no antibacterial activity on phytopathogenic bacteria *Pto* DC3000 and *Pectobacterium carotovorum* ssp. *carotovorum* 21 (*Pcc*21) (Fig. S1F). We then confirmed that valinomycin-treated plants showed bacterial growth suppression against two weak virulence pathogens (*Pto* DC3000 *Cor*^*-*^ and Δ*avrPto/*Δ*avrPtoB*) at 2- and 4-days post-inoculation (dpi) (Fig. 1A and B), while no clear difference was observed in virulent wild-type strain (Fig. S2). Considering that the AvrPto/AvrPtoB effectors and phytotoxin coronatine interfere the function of PRRs and plant hormone regulation, we hypothesized that valinomycin-induced immune enhancement is associated with PTI and hormone signaling (Shan et al., 2008; Zheng et al., 2012). This led us to further investigate putative roles of valinomycin in plant innate immunity.

**Figure 1.**
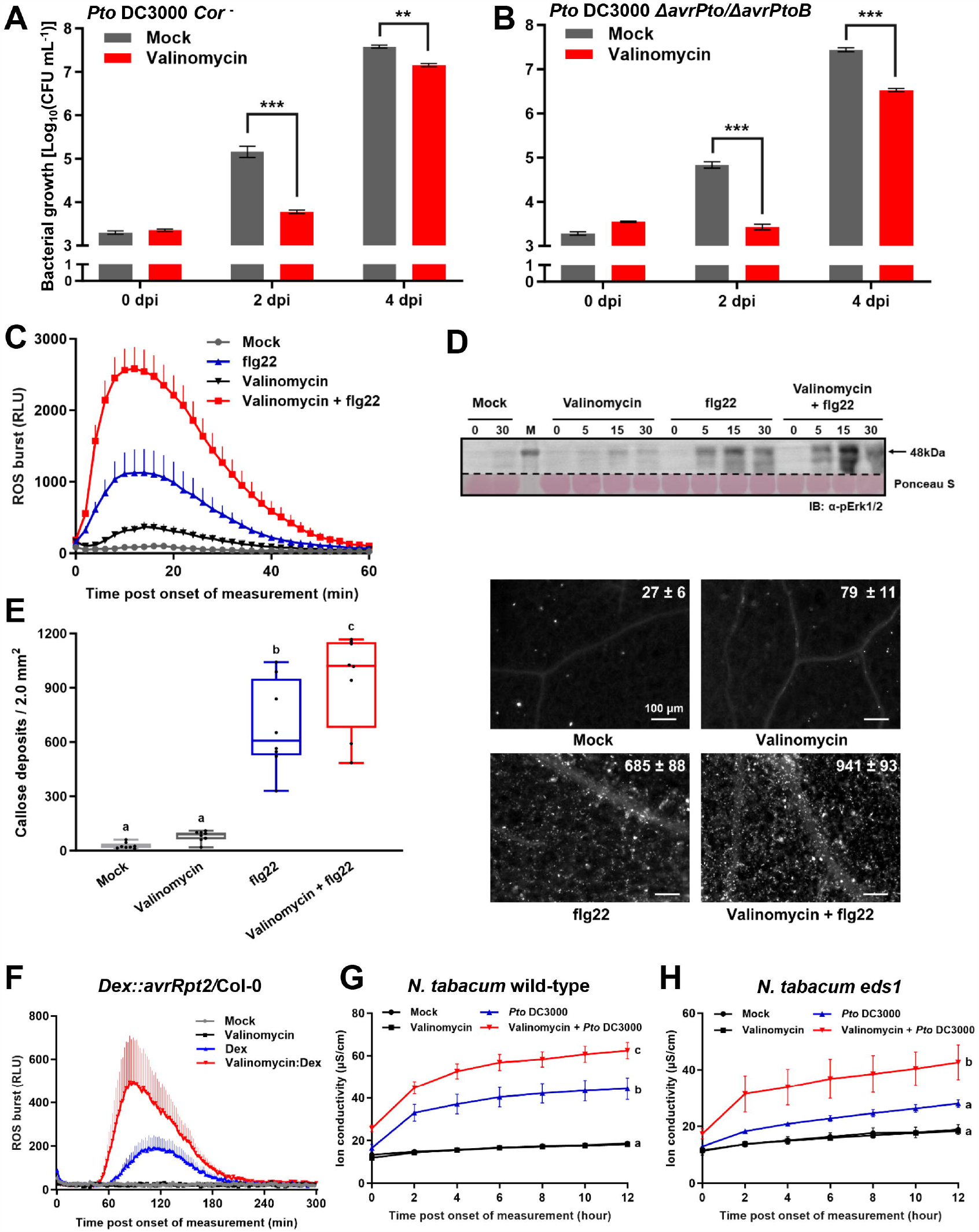
Valinomycin enhances two-layered plant innate immunity. **(A,B) Suppression of bacterial growth by valinomycin *in planta***. *Pto* DC3000 *Cor*^*-*^ mutant (A, 1 × 10^6^ CFU mL^-1^), or Δ*avrPto/*Δ*avrPtoB* mutant (B, 1 × 10^6^ CFU mL^-1^) was infiltrated on *Arabidopsis* Col-0 leaves 16 h post-valinomycin treatment. Leaves were collected on 0, 2, and 4 dpi for bacterial growth measurement (Mean ± SE, n = 3, student’s *t-*test; ** and *** indicate significance at *p* < 0.01 and *p* < 0.001 respectively, and NS indicates non-significance). **(C) Enhancement of ROS burst by valinomycin**. Twelve leaf discs of *A. thaliana* Col-0 with each assigned treatment were collected to monitor ROS burst. Luminol assay was conducted immediately after treating flg22 (Mean + SE, n = 12). **(D) Induced MAPK activation by valinomycin**. *A. thaliana* Col-0 plants pretreated with valinomycin followed by flg22 treatment were sampled at 0, 5, 15, and 30 min post-infiltration. Total protein was loaded for immunoblot with [⍰-pErk1/2 to confirm MAPK phosphorylation. Ponceau staining of RuBisCo demonstrates equal loading of each sample. **(E) Callose accumulation with valinomycin treatment**. Valinomycin and flg22 were treated as in (D). Twelve hours post-flg22 treatment, leaf discs were stained by aniline blue. Representative picture of each treatment was demonstrated along with quantification of accumulated callose (Mean ± SE, n = 10, Analysis of variance (ANOVA) followed by Duncan’s multiple range test; different letters indicate significant differences). **(F) ETI-mediated ROS burst in transgenic plant with valinomycin**. 12 leaf discs of *Dex::avrRpt2/*Col-0 were collected 16 hours post-valinomycin pretreatment. Assay was conducted immediately after treating Dex (Mean + SE, n = 12). **(G,H) Hypersensitive response with valinomycin**. *Pto* DC3000 (1 × 10^7^ CFU mL^-1^)-infected leaves of *N. tabacum* wild-type (WT) and *eds1* mutant were collected, and ion conductivity was measured for 12 h (Mean ± SE, n = 4, ANOVA followed by Duncan’s multiple range test; different letters indicate significant differences).

We addressed whether valinomycin influences on PTI outputs by employing a well-known PAMP molecule, flg22 (22-amino-acid peptide derived from *P. aeruginosa* flagellum) by monitoring valinomycin-mediated bacterial growth suppression (Fig. 1A and B). As shown in Fig. S3A, flg22-triggered ROS burst was enhanced in plants treated with various concentrations of valinomycin. Based on the MIC test result against *B. cinerea* (Fig. S1F) and ROS burst assay (Fig. S3A), we utilized 10 μM of valinomycin for further experiments. We verified that 10 μM of valinomycin treatment enhanced ROS burst triggered by flg22 (Fig. 1C). We then tested the effects of valinomycin on other PTI outputs, including MAPK activation, callose accumulation, and marker gene expression (Fig. 1D, E, and Fig. S3B, C). Consistent with ROS burst result (Fig. 1C), valinomycin-treated leaves followed by flg22 treatment displayed boosted activation of MAPK (Fig. 1D). In addition, valinomycin increased callose accumulation upon flg22 treatment compared to leaves treated with flg22 alone (Fig. 1E). Relative expression of PTI marker genes was also elevated upon the treatment of valinomycin; *FRK1* up-regulation by valinomycin pretreatment and *NHL10* up-regulation at 3 hours post-flg22 treatment (Fig. S3B, C). Together, these results provide a potential of valinomycin as a PTI-boosting natural compound.

We moved forward to test the next layer of plant immunity, ETI. To observe early response, ETI-mediated ROS burst was monitored using *Arabidopsis* transgenic plants expressing bacterial effectors (AvrRpt2) under dexamethasone (Dex)-inducible promoter. Upon Dex-treatment, valinomycin induced faster and more enhanced ROS burst compared to mock treatment (Fig. 1F). Hypersensitive response (HR), representative ETI phenotype, was monitored in tobacco plants by quantifying ion leakage from leaves. Valinomycin pretreatment enhanced HR compared to *Nicotiana benthamiana* (*N. benthamiana*) leaves infected solely with *Pto* DC3000 (Fig. S4). *Pto* DC3000 Δ*hrcC* was used as negative control that HR was dampened due to no effector delivery such as HopQ1 recognized by NLR Roq1 protein (Schultink et al., 2017). Valinomycin did not potentiate HR in response to *Pto* DC3000 *ΔhrcC* treatment, indicating that valinomycin did not activate ETI by itself (Fig. S4). We also monitored HR phenotype in *Nicotiana tabacum* (*N. tabacum*) wild-type and *eds1* mutant plants to further investigate valinomycin-boosted ETI and the involvement of salicylic acid (SA), a key defense hormone critical for HR (Venugopal et al., 2009) (Fig. 1G and 1H). The overall ion conductivity level was reduced in *eds1* mutant plants compared to wild-type. However, valinomycin displayed enhanced HR by *Pto* DC3000 in both wild-type and *eds1* mutant plants. This implies that SA is not a dominant hormone that involves in valinomycin-induced HR. In sum, we confirmed that valinomycin enhances both PTI and ETI, intensifying two-layered plant innate immunity, respectively.

### Valinomycin reinforces mutual orchestration of two layered plant immunity

To test recent important findings of mutual PTI and ETI potentiation, we measured ROS burst with flg22 and Dex-treatment in *Arabidopsis* transgenic plants expressing bacterial effectors (AvrRpm1 or AvrRpt2). The ROS burst in *Dex::avrRpt2*/Col-0 plants exhibited synergistic interaction between PTI and ETI, consistent with previous result (Fig. S5A) (Yuan et al., 2021). Moreover, we newly confirmed the mutual activation of PTI and ETI in *Dex*::*avrRpm1*/Col-0 plants (Fig. S5B). As the mutual potentiation was confirmed in both genotypes, we assessed the simultaneous immune-boosting activity on PTI and ETI byalinomycin. In both transgenic plants, valinomycin induced more robust ROS burst of PTI and ETI, strongly suggesting that valinomycin potentiated both PTI and ETI via enhancing mutual synergistic mechanism (Fig. 2A, B). We inferred that valinomycin might intensify plant immunity via SA-dependent and independent pathway based on that AvrRpm1- and AvrRpt2-mediated ETI are achieved through different signaling pathways in terms of SA-dependency (Aarts et al., 1998; Venugopal et al., 2009). Taking together, we deduced that valinomycin could boost both PTI and ETI, leading to maximized plant local innate immunity, which may function as an important signaling cue for better systemic immunity.

**Figure 2.**
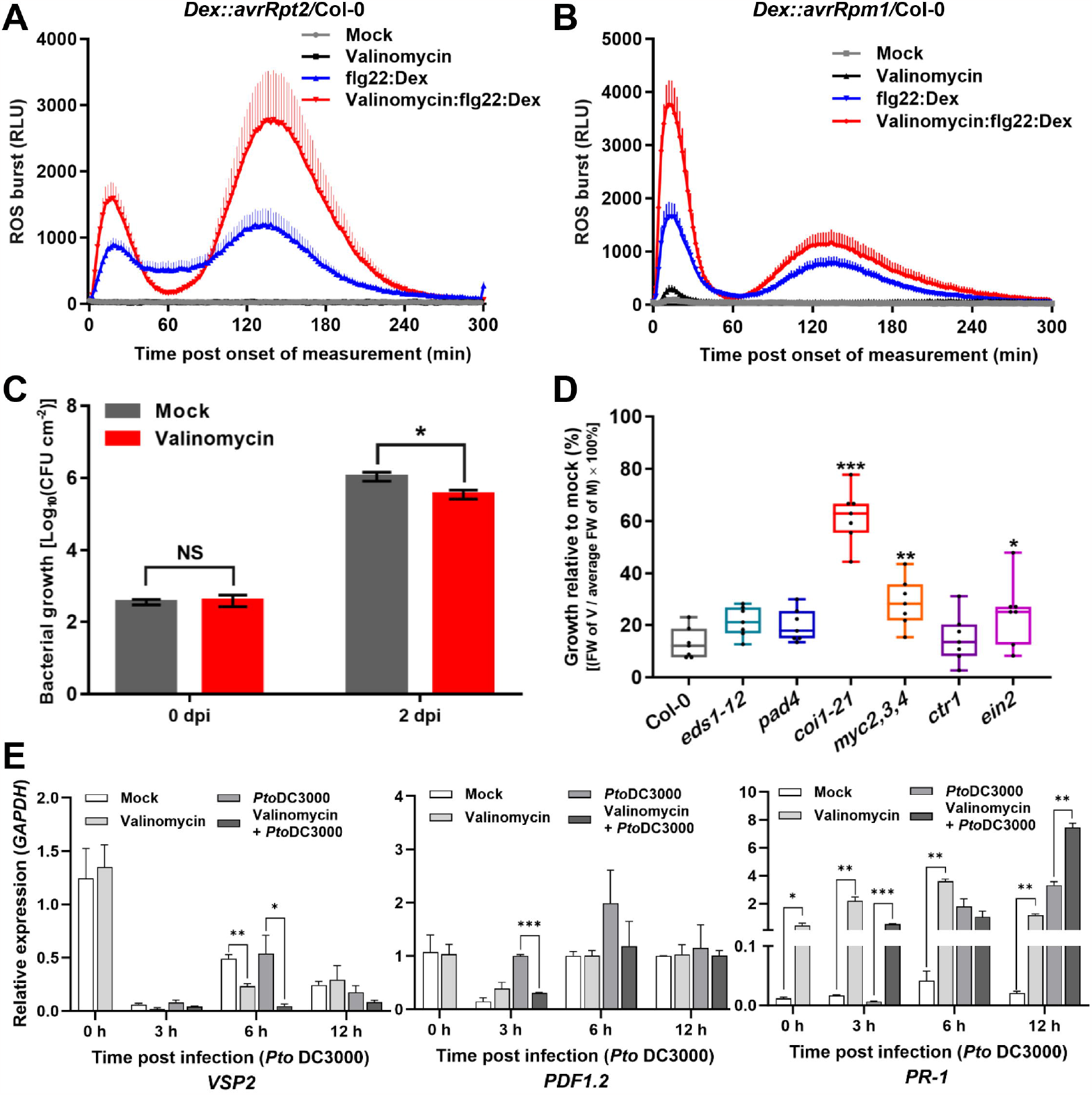
Valinomycin potentiates mutual activation of two-layered plant innate immunity and systemic immunity. **(A, B) Mutual potentiation-mediated ROS burst in transgenic plants with valinomycin**. 12 leaf discs of (A) *Dex::avrRpt2/*Col-0 and (B) *Dex::avrRpm1/*Col-0 were collected 16 hours post-valinomycin pretreatment for luminol assay. Assay was conducted immediately after treating flg22 and Dex (Mean + SE, n = 12). **(C) Systemic resistance with valinomycin**. 10 μM of valinomycin was pretreated on lower leaves 24 h prior to *Pto* DC3000 infection. *Pto* DC3000 was infiltrated on unchallenged upper leaves, and bacterial growth was observed 0 and 2 dpi (Mean ± SE, n = 3, student’s t-test; * indicates significance at p < 0.05, and NS indicates non-significance). **(D) Seedling growth inhibition with valinomycin**. Sterilized seeds were sown on 0.5X MS plates, and seedlings were transplanted into 0.5X MS liquid media with or without 10 μM of valinomycin 2 days post-germination. Seedling growth rate was measured 8 days after transplantation. Growth relative to mock was calculated by the equation [fresh weight (FW) of valinomycin-treated plant (V) / average of FW of mock-treated plants (M) × 100 (%)] (Box plots present individual replicates, n = 7, student’s *t-*test compared to Col-0; * indicates significance at *p* < 0.05). **(E) Transcriptional changes of defense signaling marker genes by valinomycin**. Leaves were sampled 0, 3, 6, 12 h after *Pto* DC3000 (10^6^ CFU mL^-1^) treatment, and relative gene expression of *VSP2* (left), *PDF1*.*2* (middle), and *PR1* (right) was analyzed by qRT-PCR. The expression of genes was normalized to the expression of house-keeping gene *GAPDH* (Mean + SE, n = 3, student’s t-test; *, **, and *** indicate significance at p < 0.05, p < 0.01, and p < 0.001 respectively).

### Valinomycin fortifies plant systemic immunity

Confirming the function of valinomycin in local plant immunity, we monitored the effect of valinomycin on systemic immunity, because local immune induction could spread systemically (Vlot et al., 2021). The bacterial growth of *Pto* DC3000 in unchallenged leaves post-valinomycin treatment at early developed leaves was significantly reduced compared to control, indicating the induction of systemic resistance (Fig. 2C). Based on this, we inferred that 1) valinomycin could translocate systemically, as other known systemic pesticides (Vryzas, 2016), 2) valinomycin-boosting local immunity shown above could trigger more robust systemic immunity, like systemic acquired resistance (SAR) or induced systemic resistance (ISR) as immune-priming (Vlot et al., 2021), 3) a combination of two scenarios could enhanced systemic immunity. In all cases, we hypothesized that valinomycin could fortify systemic immunity, working as an immune-priming agent or a mobile signaling molecule, even though further investigation is necessary to determine fine-tuned mechanisms (Jung et al., 2009).

Considering plant immune-priming such as SAR and ISR, we focused on plant hormones required for systemic immunity (Pieterse et al., 2009). First, seedling growth inhibition assay of Col-0 and plant hormone mutant plants (*eds1-12, pad4, coi1-21, myc2,3,4, ctr1*, and *ein2-1*) was performed (Bredow et al., 2019). Overall, valinomycin reduced fresh weight of plants in all genotypes, suggesting that no dominant hormonal pathway be involved in the activity of valinomycin (Fig. 2D). Nevertheless, the seedling growth of jasmonic acid (JA) signaling mutant *coi1-21* demonstrated minimum defect by valinomycin treatment. Another JA signaling mutant *myc2,3,4* and ethylene (ET) signaling mutant *ein2-1* displayed less growth defect following *coi1-21* (Fig. 2D). This implies that valinomycin-induced immune-boosting is more dependent on JA and ET than SA (Fig. 2A, B). Interestingly, relative gene expression of *VSP2* (JA signaling-related gene) and *PDF1*.*2* (JA/ET-responsive defense gene) was downregulated post-*Pto* DC3000 infection (Fig. 2E). In contrast, *PR-1* (SA-responsive defense marker gene) was upregulated highly by valinomycin (Fig. 2E).

Considering antagonistic interaction between SA and JA and bacterial growth suppression phenotype in Figure 1A and B combined with seedling growth suppression (Fig. 2B) and qRT results (Fig. 2E), we speculate that valinomycin may reduce JA signaling upon treatment, leading to gradual upregulation of SA for successful enhancement of local and systemic immune responses (Pieterse et al., 2009). Hemibiotroph *Pto* DC3000 secretes coronatine, JA-isoleucine mimic, to increase virulence by activating JA signaling that results in inhibition of SA accumulation (Mur et al., 2006; Zheng et al., 2012). Thus, valinomycin may counteract on the infection strategy of *Pto* DC3000 through attenuating JA-responsive signaling, leading to more SA-related immune responses.

Our results demonstrate that the local immune-boosting activity of valinomycin may be dependent on JA and/or ET signaling pathway based on seedling growth assay result (Fig. 2D, E). Moreover, bacterial growth suppression assay with *Pto* DC3000 *Cor*^*-*^ and *Pto* DC3000 Δ*avrPto/*Δ*avrPtoB* strongly support the compromised virulence activity via JA and increased disease resistance by SA, respectively (Fig. 1A, B) (Tateda et al., 2014; Zheng et al., 2012). Thus, valinomycin-induced local immune-boosting is likely to be dependent on JA at the early pathogen infection and enhanced SA-dependent immune response later, which eventually can enhance systemic immunity by priming.

In conclusion, this study highlights the potential of valinomycin as a biocontrol agent with dual role in antifungal and plant immune-boosting activity. Multi-functional biocontrol agents are efficient in controlling complex pathogen infections occurred naturally, reducing the possibility of antibiotic resistance of pathogens. Most importantly, we uncovered the role of valinomycin in two tiers of plant innate immunity thoroughly, including intensified PTI, ETI, and PTI-ETI mutual potentiation, mainly focusing on overall disease suppression phenotype. Valinomycin-triggered systemic immunity was observed as well, suggesting this natural compound as a putative broad-spectrum plant immune-priming biocontrol agent (Fig. S6). While further investigation of the detailed mechanism is warranted, our findings expand the potential utility of natural compounds as promising alternatives that could replace or complement the need for antibiotics or chemical treatment to control plant disease in agriculture.

## ACKNOWLEDGEMENTS

We express our gratitude to Prof. Eunkyoo Oh at Korea University for generously providing *myc 2,3,4* triple mutant utilized in this study. We also extend our appreciation to Prof. Sang Hee Kim at Gyeongsang National University for his meticulous review of the manuscript. This study was supported by the Korea University Grant (K239821) and the Rural Development Agent (PJ015871032021).

## REFERENCES

Aarts, N., Metz, M., Holub, E., Staskawicz, B. J., Daniels, M. J., & Parker, J. E. (1998). Different requirements for EDS1 and NDR1 by disease resistance genes define at least two R gene-mediated signaling pathways in Arabidopsis. Proceedings of the national academy of sciences, 95(17), 10306–10311.

Bredow, M., Sementchoukova, I., Siegel, K., & Monaghan, J. (2019). Pattern-triggered oxidative burst and seedling growth inhibition assays in Arabidopsis thaliana. JoVE (Journal of Visualized Experiments)(147), e5.9437.

Chang, M., Chen, H., Liu, F., & Fu, Z. Q. (2022). PTI and ETI: convergent pathways with diverse elicitors. Trends in plant science, 27(2), 113–115.

Cohen, Y., Vaknin, M., & Mauch-Mani, B. (2016). BABA-induced resistance: milestones along a 55-year journey. Phytoparasitica, 44, 513–538.

Cooper, O., Seo, H., Andrabi, S., Guardia-Laguarta, C., Graziotto, J., Sundberg, M., McLean, J. R., Carrillo-Reid, L., Xie, Z., Osborn, T., Hargus, G., Deleidi, M., Lawson, T., Bogetofte, H., Perez-Torres, E., Clark, L., Moskowitz, C., Mazzulli, J., Chen, L., … Isacson, O. (2012). Pharmacological Rescue of Mitochondrial Deficits in iPSC-Derived Neural Cells from Patients with Familial Parkinson’s Disease. Science Translational Medicine, 4(141), 141ra190–141ra190.

Dickey, S. W., Cheung, G. Y., & Otto, M. (2017). Different drugs for bad bugs: antivirulence strategies in the age of antibiotic resistance. Nature Reviews Drug Discovery, 16(7), 457–471.

Dodds, P. N., & Rathjen, J. P. (2010). Plant immunity: towards an integrated view of plant–pathogen interactions. Nature Reviews Genetics, 11(8), 539–548.

Farha, M. A., Verschoor, C. P., Bowdish, D., & Brown, E. D. (2013). Collapsing the proton motive force to identify synergistic combinations against Staphylococcus aureus. Chemistry & biology, 20(9), 1168–1178.

Jones, J. D., & Dangl, J. L. (2006). The plant immune system. Nature, 444(7117), 323–329.

Jung, H. W., Tschaplinski, T. J., Wang, L., Glazebrook, J., & Greenberg, J. T. (2009). Priming in systemic plant immunity. Science, 324(5923), 89–91.

Kang, J. E., Hwang, S., Yoo, N., Kim, B. S., & Chung, E.-H. (2022). A resveratrol oligomer, hopeaphenol suppresses virulence activity of Pectobacterium atrosepticum via the modulation of the master regulator, FlhDC. Frontiers in Microbiology, 4103.

Kang, J. E., Yoo, N., Jeon, B. J., Kim, B. S., & Chung, E.-H. (2022). Resveratrol oligomers, a plant-driven natural product with anti-virulence and plant immune-priming roles. Frontiers in plant science, 1553.

Mur, L. A., Kenton, P., Atzorn, R., Miersch, O., & Wasternack, C. (2006). The outcomes of concentration-specific interactions between salicylate and jasmonate signaling include synergy, antagonism, and oxidative stress leading to cell death. Plant physiology, 140(1), 249–262.

Park, C.-N., Lee, J.-M., Lee, D.-H., & Kim, B.-S. (2008). Antifungal activity of valinomycin, a peptide antibiotic produced by Streptomyces sp. strain M10 antagonistic to Botrytis cinerea. Journal of microbiology and biotechnology, 18(5), 880–884.

Pieterse, C. M., Leon-Reyes, A., Van der Ent, S., & Van Wees, S. C. (2009). Networking by small-molecule hormones in plant immunity. Nature chemical biology, 5(5), 308–316.

Pruitt, R. N., Gust, A. A., & Nürnberger, T. (2021). Plant immunity unified. Nature plants, 7(4), 382–383.

Schultink, A., Qi, T., Lee, A., Steinbrenner, A. D., & Staskawicz, B. (2017). Roq1 mediates recognition of the Xanthomonas and Pseudomonas effector proteins XopQ and HopQ1. The Plant Journal, 92(5), 787–795.

Shan, L., He, P., Li, J., Heese, A., Peck, S. C., Nürnberger, T., Martin, G. B., & Sheen, J. (2008). Bacterial effectors target the common signaling partner BAK1 to disrupt multiple MAMP receptor-signaling complexes and impede plant immunity. Cell host & microbe, 4(1), 17–27.

Silva, L. N., Zimmer, K. R., Macedo, A. J., & Trentin, D. S. (2016). Plant natural products targeting bacterial virulence factors. Chemical reviews, 116(16), 9162–9236.

Tateda, C., Zhang, Z., Shrestha, J., Jelenska, J., Chinchilla, D., & Greenberg, J. T. (2014). Salicylic acid regulates Arabidopsis microbial pattern receptor kinase levels and signaling. The Plant Cell, 26(10), 4171–4187.

Venugopal, S. C., Jeong, R.-D., Mandal, M. K., Zhu, S., Chandra-Shekara, A., Xia, Y., Hersh, M., Stromberg, A. J., Navarre, D., & Kachroo, A. (2009). Enhanced disease susceptibility 1 and salicylic acid act redundantly to regulate resistance gene-mediated signaling. PLoS genetics, 5(7), e1000545.

Vlot, A. C., Sales, J. H., Lenk, M., Bauer, K., Brambilla, A., Sommer, A., Chen, Y., Wenig, M., & Nayem, S. (2021). Systemic propagation of immunity in plants. New Phytologist, 229(3), 1234–1250.

Vryzas, Z. (2016). The plant as metaorganism and research on next-generation systemic pesticides–prospects and challenges. Frontiers in Microbiology, 7, 1968.

Yuan, M., Jiang, Z., Bi, G., Nomura, K., Liu, M., Wang, Y., Cai, B., Zhou, J.-M., He, S. Y., & Xin, X.-F. (2021). Pattern-recognition receptors are required for NLR-mediated plant immunity. Nature, 592(7852), 105–109.

Zheng, X.-y., Spivey, N. W., Zeng, W., Liu, P.-P., Fu, Z. Q., Klessig, D. F., He, S. Y., & Dong, X. (2012). Coronatine promotes Pseudomonas syringae virulence in plants by activating a signaling cascade that inhibits salicylic acid accumulation. Cell host & microbe, 11(6), 587–596.

Zipfel, C., Robatzek, S., Navarro, L., Oakeley, E. J., Jones, J. D., Felix, G., & Boller, T. (2004). Bacterial disease resistance in Arabidopsis through flagellin perception. Nature, 428(6984), 764–767.

