## Supporting information for "A natural cyclic peptide valinomycin enhances plant innate immunity"

### SUPPLEMENTARY MATERIALS AND METHODS

#### Plant materials and growth conditions

*A. thaliana* Col-0 accessions were used for *in planta* experiments. Col-0, isogenic mutants, and transgenic plants were sown in a potting mix with 3:1:1 (v/v) ratio of Sungro mix #4 (Sun Gro Horticulture, USA) : vermiculite : perlite supplemented with 250 PPM Peter's special 20-20-20 (w/w) fertilizer (Everris NA, The Netherlands). Seeds were sterilized before sowing by chlorine gas (30 mL of NaOCl supplemented with 3 mL of 6 N HCl) or by 50% NaOCl. Seeds sowed on soil or MS media kept in the dark at 4 °C for 3 days for vernalization. Plants were grown in a growth chamber (Hanbaek scientific co., Republic of Korea) under 9 h light/15 h dark cycle (21 °C day/18 °C night, 60% relative humidity (RH)). Two-week-old seedlings were transplanted, and four or five-week-old plants were used for the following experiments.

*N. benthamiana* and *N. tabacum* seeds were sown in the same potting mix and were grown in growth rooms 12 h light/12 h dark cycle (24 °C, 60% RH). After two weeks, the tobacco plants were transplanted, and five-week-old plants were used for the following experiments.

#### Anti-microbial activity

Paper disc bioassay was performed as described in Kim et al. (2020). The conidia of *B. cinerea* were harvested using sterile distilled water and then filtered through sterile miracloth to remove mycelial fragments from the suspension. The number of conidia in the suspension was determined by direct microscopic counting in a hemocytometer. Paper discs saturated with antifungal metabolites were loaded onto potato dextrose agar (PDA) medium (0.7% agar) supplemented with the conidial suspension of *B. cinerea* ($1 \times 10^6$ conidia $mL^{-1}$). The plates were incubated at 22 °C until a visually clear inhibition zone appeared around the discs.

Minimal inhibitory concentration (MIC) assay of valinomycin was performed against *B. cinerea* B05.10, *Pto* DC3000, and *Pcc*21, as described in Kang et al. (2020) with minor modifications. For *B. cinerea*, serial ten-fold dilutions of valinomycin ranging from 10 nM to 100 μM with the conidial suspension ($1 \times 10^4$ conidia $mL^{-1}$) were used to determine MIC in PDB medium in 96-well plate. The plate was incubated at 22 °C for a week. For bacterial pathogens, overnight cultures of *Pto* DC3000 and *Pcc*21 were diluted to $1 \times 10^6$ colony-forming unit (CFU) $mL^{-1}$ respectively with King's B (KB) medium and lysogeny broth (LB) medium and incubated with the same concentrations of valinomycin as above at 28 °C for four days.

### Isolation and identification of active compound valinomycin

A single colony of S8 strain grown on tryptic soy agar (TSA) was transferred into a 1-L baffled flask containing 100 mL of tryptic soy broth (TSB), and cultured using a rotary shaker at 200 rpm at 28 °C. After 3 days, 1% of the seed culture was used as an inoculum for 4 L scale culture. After 5 days of incubation at 200 rpm at 28 °C, the culture was centrifuged at 10,000 × g for 10 min to separate bacterial cells and cell-free supernatant. The cells were extracted with methanol and the supernatant was extracted with an equal volume of ethyl acetate and butanol in order. Each extract was concentrated *in vacuo* on a rotary evaporator, and dissolved in 8 mL of a high-performance liquid chromatography (HPLC) grade methanol. 10 mL of culture volume of each extract was loaded onto the paper disc to determine antifungal activity against *B. cinerea* B05.10.

To purify active ingredients from the concentrated extract, a reverse phase chromatography was performed using Sep-Pak C18 cartridge (Waters, USA). Elution was performed with stepwise gradients of water and methanol (v/v; 100:0, 80:20, 60:40, 40:60, 20:80, and 0:100). Each fraction was tested for its antifungal activity through paper disc assay. The active fraction was subjected to additional purification step using HPLC with C18 reverse-phase column (25 cm × 10 mm, S-5 μm) (Supelco, USA) and pentafluorophenyl (PFP) column (150 mm × 4.6 mm, S-4 μm) (YMC, Japan). Chromatography was performed using a linear gradient dilution system (20-95% methanol in $H_2O$) for 20 min, followed by isocratic elution of 95% aqueous methanol for 20 min at a flow rate of 2 mL $min^{-1}$ for C18 column and 1 mL $min^{-1}$ for PFP column. HPLC chromatograms were monitored at an absorbance of 220 nm, and the active compound was obtained as a white powder (Fig. S1A-C). Electrospray ionization high-resolution mass spectrometry (ESI-HRMS) was performed to identify the active compound purified from the culture of *Streptomyces bacillaris* sp. S8 (Fig. S1D). After mass spectrometry analysis, the $^{1}H$-nuclear magnetic resonance (NMR) spectra of the compound were recorded using a Varian 500 NMR spectrometer (Varian, USA) at room temperature. The $^{1}H$-NMR (500 MHz) spectrum was measured in chloroform-*d*. Based on the spectroscopic data (Fig. S1E), the structure of the active compound was determined (Ye et al., 2017).

### Chemical treatment

Valinomycin dissolved in methanol (1 mM) was diluted 100 times with dd$H_2O$ for hand-infiltration in leaves. For mock treatment, same volume of HPLC grade methanol was added to dd$H_2O$ (1% methanol). Plants were pretreated with 10 μM of valinomycin and 1% methanol as a mock before pathogen or flg22 treatment.

### ROS burst measurement

To measure the amount of ROS production, chemiluminescence assay was performed following description of Smith and Heese (2014) with minor modifications. 5-week-old *A. thaliana* Col-0 plants were treated with 10 μM of valinomycin or 1% methanol (mock) by hand infiltration. After 16 hours, leaf discs were placed in the white 96-well plate containing 200 μL of sterile water in each well. The plate was kept under the light for three hours to minimize wound response. After gently removing the water, 100 μL of reaction solution containing 100 nM of flg22 obtained from *P. aeruginosa*, 30 μg $mL^{-1}$ of luminol, and 20 μg $mL^{-1}$ of horseradish peroxidase (HRP) were added to each well. Luminescence of the reaction was detected every 2 min for 60 min by using Centro luminometer (Berthold technologies, Germany).

The ROS burst triggered by PTI and ETI was confirmed in Dex::*avrRpt2*/Col-0 and Dex::*avrRpm1*/Col-0 plants, respectively harboring *avrRpt2* and *avrRpm1* genes under the control of Dexamethasone (Dex)-inducible promoter. Valinomycin treatment and leaf disc preparation were fulfilled as mentioned above with a reaction solution consisting of 100 nM of flg22, 5 μM of Dex, 30 μg $mL^{-1}$ of luminol, and 20 μg $mL^{-1}$ of HRP. Chemiluminescence analysis was conducted every 2 min for 5 hours.

### Protein extraction and immunoblot analysis

MAPK activation was observed as described in Chung et al. (2014). *A. thaliana* was treated with 10 μM of valinomycin or 1% methanol (mock) 16 hours prior to treatment of 100 nM of flg22. To compare MAPK activities in the plants treated with valinomycin or 1% methanol, two leaves were collected at each time point (0, 5, 15, 30 min after flg22 treatment) and ground after freezing with liquid nitrogen utilizing tissue homogenizer (Benchmark scientific, USA) (4,000 rpm, 30 sec, 2 cycles). A total protein of Col-0 leaves was extracted using protein extraction buffer containing 100 mM of Tris pH 8.0, 1% Triton X 100, 0.1% SDS, 10 mM of Dithiothreitol, 1X protease inhibitor cocktail, and 1X phosphatase inhibitor cocktail. The lysates were centrifuged at 14,000 rpm at 4 °C for 10 min. Supernatant equivalent to 500 ng of total protein was electrophoresed through 10% SDS-PAGE, followed by semi-dry transfer onto PVDF membrane. Immunoblots were performed with a Phospho-p44/42 MAPK antibody (Erk1/2: Cell Signaling Technology, USA, 1:1,000 dilution) and then HRP-conjugated secondary antibody (Santa Cruz Biotechnology, USA, 1:5,000 dilution) was added. The blots were developed by enhanced chemiluminescence (ECL) (GenDEPOT, USA) and the images of the immunoblots were captured in a Chemidoc (Bio-Rad, USA).

### Callose quantification

Callose quantification was performed following the description mentioned in Chung et al. (2014) with minor modifications. 5-week-old Col-0 plants were treated with 10 μM valinomycin or 1% methanol by hand-infiltration with 1 mL needleless syringe. After 16 hours, 100 nM of flg22 was infiltrated into the leaves pretreated with valinomycin or 1% methanol. The plants were kept in the normal growth condition (under 9 h light/15 h dark cycle, 60% RH) overnight, and leaf discs were collected 12 hours post-treatment. The discs were placed into 15 mL conical tube containing 100% ethanol to draw out chlorophyll at 37 °C overnight and then washed with water several times. After completely removing ethanol, 0.01% (w/v) aniline blue solution (dissolved in 150 mM of $K_2HPO_4$, pH 9.5) was added to stain the callose. Staining was performed at room temperature overnight covered with aluminum foil to protect the dye from light. Stained leaves were washed gently with distilled water 3 times. 17 μL of 40% glycerol solution was loaded onto a slide glass and the leaf discs were placed on the slide glass. Callose deposits were visualized by fluorescent microscope with 100 x magnification (ZEISS, Germany), and the images were captured by the mounted camera. ImageJ software (National Institutes of Health, USA) was used to quantify callose deposits from the images.

### Conductivity analysis for HR quantification

Bacterial suspension of *Pto* DC3000 wild type and *ΔhrcC* mutant strain ($1 \times 10^7$ CFU $mL^{-1}$) was infiltrated into leaves of 5-week-old *N. benthamiana* and *N. tabacum* plants 16 h after treatment with valinomycin or 1% methanol. Four leaf discs from each treatment with 4 biological replicates were submerged into clear glass tubes containing 6 mL of distilled water 12 h post-infection. Ion leakage was immediately measured by using a conductivity meter (Apera instruments, USA) at room temperature, every 2 h for 12 hours in total.

For phenotype analysis of HR, *Pto* DC3000 ($1 \times 10^6$ CFU $mL^{-1}$) was inoculated on *N. benthamiana*. One day post-*Pto* DC3000 treatment, picture of inoculated leaf was taken by Chemidoc (Bio-Rad, USA).

### Bacterial growth suppression assay

The growth of *Pto* DC3000 wild-type, C*or*$^-$, and *ΔavrPto/ΔavrPtoB* in Col-0 was confirmed as described in Tornero and Dangl (2001). Col-0 was treated with 10 μM of valinomycin or 1% methanol 16 hours prior to *Pto* DC3000 infection. Bacterial suspension was adjusted to $10^5$ CFU

$mL^{-1}$ for wild-type and $10^6$ CFU $mL^{-1}$ for *Cor*$^-$ and *ΔavrPto/ΔavrPtoB* to infect the plants. Bacterial growth was determined 0, 2, and 4 days post-infection (dpi) of bacterial suspension.

Systemic resistance in *A. thaliana* was confirmed as described in Rufián et al. (2019) with minor modifications. 5-week-old *A. thaliana* Col-0 plants with about 12 rosette leaves were treated with 10 μM of valinomycin or 1% methanol on 3 lower rosette leaves. *Pto* DC3000 ($1 \times 10^5$ CFU $mL^{-1}$) was inoculated on upper unchallenged rosette leaves 24 hours post-valinomycin treatment. The bacterial growth was measured at 0 and 2 dpi from *Pto* DC3000-treated leaves without chemical treatment.

**Seedling growth inhibition assay**

The growth of *A. thaliana* was monitored as following description of Bredow et al. (2019). *A. thaliana* Col-0 wild-type and mutant seeds (*eds1-12, pad4, coi1-21, myc2,3,4, ctr1, ein2-1*) were sown in 0.5 x Murashige and Skoog (MS) (Duchefa biochemie, Netherlands) plates. After 3 days of vernalization at 4 °C, seeds were germinated in the growth chamber. After four days, the seedlings were transplanted into 48-well plate each containing 1 mL of 0.5X MS liquid media (supplemented with 1% sucrose) with 10 μM of valinomycin or 1% methanol. Seedling growth was monitored every day, and fresh weight (FW) was measured on Day 8. The growth relative to mock (%) was calculated by the equation [FW of valinomycin-treated plant / mean of FW of mock-treated plants] × 100 (%). Relative growth was calculated from the means of seven replicates.

**qRT-PCR analysis for marker gene expression**

5-week-old *A. thaliana* Col-0 plants were pretreated with 10 μM valinomycin by hand-infiltration. After 16 h, *Pto* DC3000 ($1 \times 10^6$ CFU $mL^{-1}$) was inoculated with 1-mL syringe without needle, and sample was collected after 0, 3, 6, and 12 h post-infection, and frozen immediately in liquid nitrogen. RNA extraction of collected leaves was performed as described previously (Kang et al., 2016) with minor modifications. RNA was extracted using RNeasy kit (Qiagen, Germany), and 500 ng of extracted RNA was converted to DNA using cDNA synthesis kit (Bio-Rad, USA). qRT-PCR was performed with the primers listed in Table S1, and amplification was conducted utilizing QuantStudio™ 3 real-time PCR system (Applied Biosystems, USA) following manufacturer's protocol. GAPDH gene was used as an internal standard, and normalized gene expression was calculated by comparative $C_T$ method ($2^{\Delta Ct(\text{target gene})}/2^{\Delta Ct(\text{internal control gene})}$) as described in Kang et al. (2022).

### Statistical analysis

The Statistical Analysis System software (SAS Institute, USA) was used to analyze the data. Student's *t*-test ($P < 0.05$) was conducted to determine statistical significance between valinomycin and mock-treated plant. One-way analysis of variance (ANOVA) was performed to compare the amount of callose deposition and intensity of ion conductivity between different treatments. Mean separation was conducted using Duncan's multiple range test at $\alpha = 0.05$.

### Graphical image generation

The working model was generated by BioRender with publication permission.

**Supplemental References**


Bredow, M., Sementchoukova, I., Siegel, K., & Monaghan, J. (2019). Pattern-triggered oxidative burst and seedling growth inhibition assays in *Arabidopsis thaliana*. *JoVE (Journal of Visualized Experiments)*(147), e59437.

Chung, E.-H., El-Kasmi, F., He, Y., Loehr, A., & Dangl, J. L. (2014). A plant phosphoswitch platform repeatedly targeted by type III effector proteins regulates the output of both tiers of plant immune receptors. *Cell host & microbe*, *16*(4), 484-494.

Kang, J. E., Han, J. W., Jeon, B. J., & Kim, B. S. (2016). Efficacies of quorum sensing inhibitors, piericidin A and glucopiericidin A, produced by *Streptomyces xanthocidicus* KPP01532 for the control of potato soft rot caused by *Erwinia carotovora* subsp. *atroseptica*. *Microbiological research*, *184*, 32-41.

Kang, J. E., Hwang, S., Yoo, N., Kim, B. S., & Chung, E.-H. (2022). A resveratrol oligomer, hopeaphenol suppresses virulence activity of *Pectobacterium atrosepticum* via the modulation of the master regulator, FlhDC. *Frontiers in Microbiology*, 4103.

Kang, J. E., Jeon, B. J., Park, M. Y., Yang, H. J., Kwon, J., Lee, D. H., & Kim, B. S. (2020). Inhibition of the type III secretion system of *Pseudomonas syringae* pv. *tomato* DC3000 by resveratrol oligomers identified in *Vitis vinifera* L. *Pest management science*, *76*(7), 2294-2303.

Kim, Y. H., Kim, G. H., Yoon, K. S., Shankar, S., & Rhim, J.-W. (2020). Comparative antibacterial and antifungal activities of sulfur nanoparticles capped with chitosan. *Microbial Pathogenesis*, *144*, 104178.

Rufián, J. S., Rueda-Blanco, J., Beuzón, C. R., & Ruiz-Albert, J. (2019). Protocol: an improved method to quantify activation of systemic acquired resistance (SAR). *Plant methods*, *15*, 1-8.

Smith, J. M., & Heese, A. (2014). Rapid bioassay to measure early reactive oxygen species production in *Arabidopsis* leave tissue in response to living *Pseudomonas syringae*. *Plant methods*, *10*(1), 1-9.

Tornero, P., & Dangl, J. L. (2001). A high-throughput method for quantifying growth of phytopathogenic bacteria in *Arabidopsis thaliana*. *The Plant Journal*, *28*(4), 475-481.

Ye, X., Anjum, K., Song, T., Wang, W., Liang, Y., Chen, M., Huang, H., Lian, X.-Y., & Zhang, Z. (2017). Antiproliferative cyclodepsipeptides from the marine actinomycete *Streptomyces* sp. P11-23B downregulating the tumor metabolic enzymes of glycolysis, glutaminolysis, and lipogenesis. *Phytochemistry*, *135*, 151-159.

### SUPPLEMENTARY TABLE

#### Table S1. Primers used in this study

| Primer | Sequence |
|---|---|
| F_*GAPDH* | 5'-CTCCCATGTTTGTTGTTGGTGTCA-3' |
| R_*GAPDH* | 5'-CTTCCACCTCTCCAGTCCTTCATT-3' |
| F_*VSP2* | 5'-ATGCCAAAGGACTTGCCCTA-3' |
| R_*VSP2* | 5'-CGGGTCGGTCTTCTCTGTTC-3' |
| F_*PDF1.2* | 5'-CGAGAAGCCAAGTGGGACAT-3' |
| R_*PDF1.2* | 5'-ACTTGTGTGCTGGGAAGACA-3' |
| F_*PR-1* | 5'-ACGGGGAAAACTTAGCCTGG-3' |
| R_*PR-1* | 5'-CACGTGGCAGAAGTTGTGTG-3' |
| F_*FRK1* | 5'-ATAGTTCACAGAGATGTGAAG-3' |
| R_*FRK1* | 5'-AGTCGAATAGTACTCGGGGTCA-3' |
| F_*NHL10* | 5'-TTCCTGTCCGTAACCCAAAC-3' |
| R_*NHL10* | 5'-CCCTCGTAGTAGGCATGAGC-3' |

**SUPPLEMENTARY FIGURES**

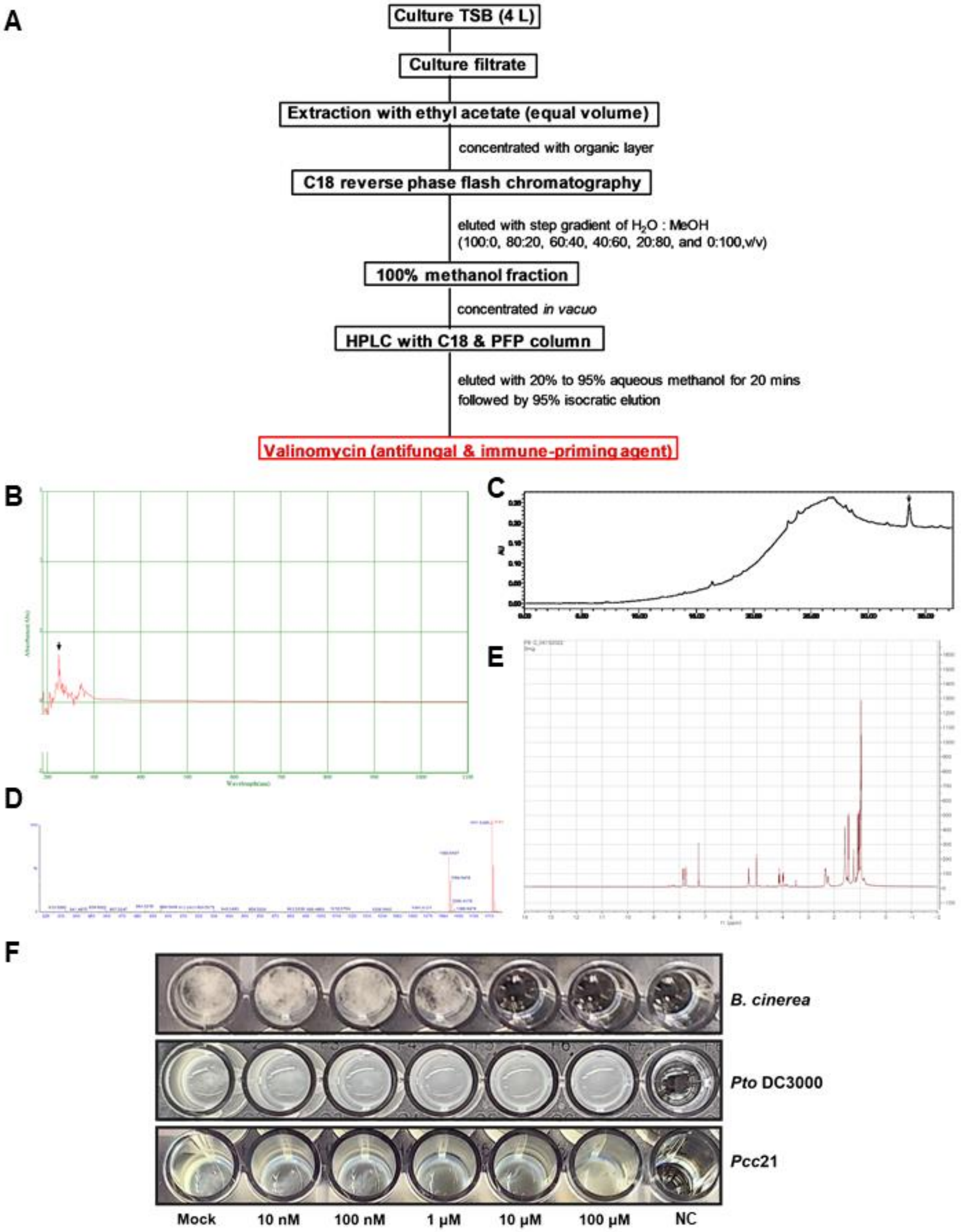

**Figure S1. Streptomyces-derived valinomycin possesses antifungal activity against *B. cinerea,* but no antibacterial activity against bacterial phytopathogens.**

**(A)** Antifungal activity-guided purification procedure of valinomycin produced by *Streptomyces bacillaris* sp. S8.

**(B)** UV scan profile of valinomycin. Valinomycin exhibited maximum absorbance at 220 nm, which is pointed by the arrow.

**(C)** HPLC profile of 100% methanol fraction. The eluate was monitored at 220nm, and chromatography was conducted using a linear gradient dilution system (20-95% methanol in $H_2O$) for 20 min, followed by isocratic elution of 95% aqueous methanol for 20 min. The peak pointed by the arrow indicates valinomycin.

**(D)** Electrospray ionization high-resolution mass spectrometry (ESI-HRMS) data of valinomycin.

**(E)** $^1$H-NMR data of valinomycin.

**(F)** Minimum Inhibitory Concentration (MIC) was tested against *B. cinerea*, *Pt*o DC3000, and *Pcc*21. Valinomycin was added to each well of the 96-well plate to meet 10 nM, 100 nM, 1 µM, 10 µM, and 100 µM as a final concentration. Mock contained bacterial/fungal suspension without valinomycin, and negative control (NC) contained only the medium that each microbial strain was cultured in. The suspension of *B. cinerea* was diluted to $10^4$ conidia $mL^{-1}$ with PDB medium and grown for 7 days. *Pto* DC3000 was diluted to $10^6$ CFU $mL^{-1}$ with KB medium and grown for 4 days. *Pcc*21 was diluted to $10^6$ CFU $mL^{-1}$ with LB medium and grown for 4 days.

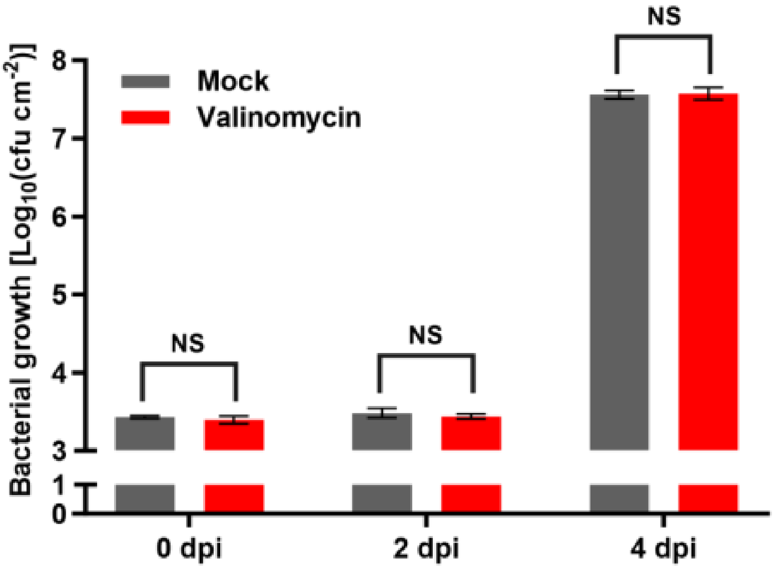


**Figure S2. Valinomycin does not suppress bacterial growth of *Pto* DC3000 wild-type strain in Arabidopsis.** Bacterial suspension of *Pto* DC3000 ($1 \times 10^5$ CFU $mL^{-1}$) was infiltrated on *A. thaliana* Col-0 leaves 16 h post-valinomycin treatment. Leaves were collected on 0- and 2-days post-infection (dpi) for bacterial growth measurement (Mean ± SE, n = 3, student's *t*-test; NS indicates non-significance).

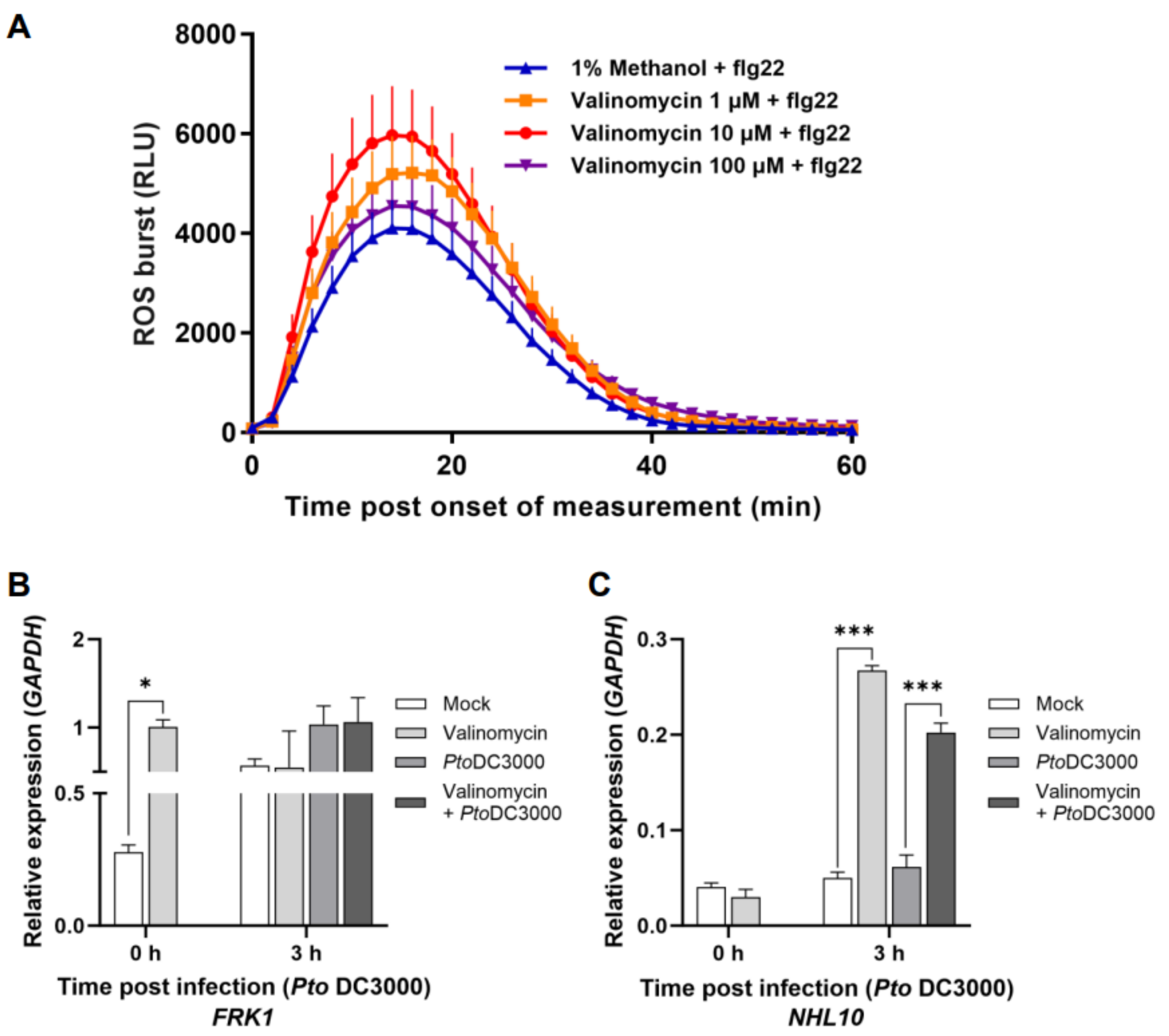


**Figure S3. Valinomycin boosts PTI responses by inducing expression of PTI maker genes.**

**(A)** ROS burst assay with different concentrations of valinomycin**.** Twelve leaf discs of *A. thaliana* Col-0 with each assigned treatment were collected for luminol assay to monitor ROS burst (Mean + SE, n = 12).

**(B, C)** Transcriptional changes in PTI signaling marker genes**.** Expression of (B) *FRK1* and (C) *NHL10* was monitored 0 and 3 h post-*Pto* DC3000 infection by quantitative polymerase chain reaction. Relative gene expression was normalized to the expression of house-keeping gene *GAPDH*, and error bars indicate SE (n = 3). Student's *t*-test was conducted to determine the statistical difference between mock-treated and valinomycin-treated samples. * and *** indicate significance at $p < 0.05$ and $p < 0.001$ respectively.

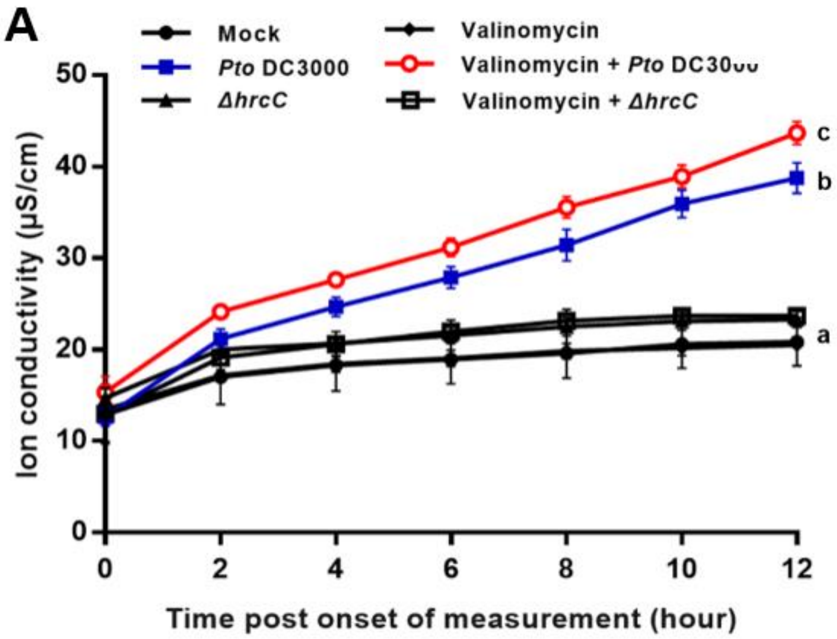


**Figure S4. Valinomycin enhances HR responses in *N. benthamiana*..** *Pto* DC3000 ($1 \times 10^7$ CFU $mL^{-1}$) was inoculated on *N. benthamiana* leaves 16 h post-valinomycin treatment. Infected leaves were collected 12 h post-treatment, and ion conductivity was measured for 12 h (ANOVA followed by Duncan's multiple range test; different letters indicate significant differences).

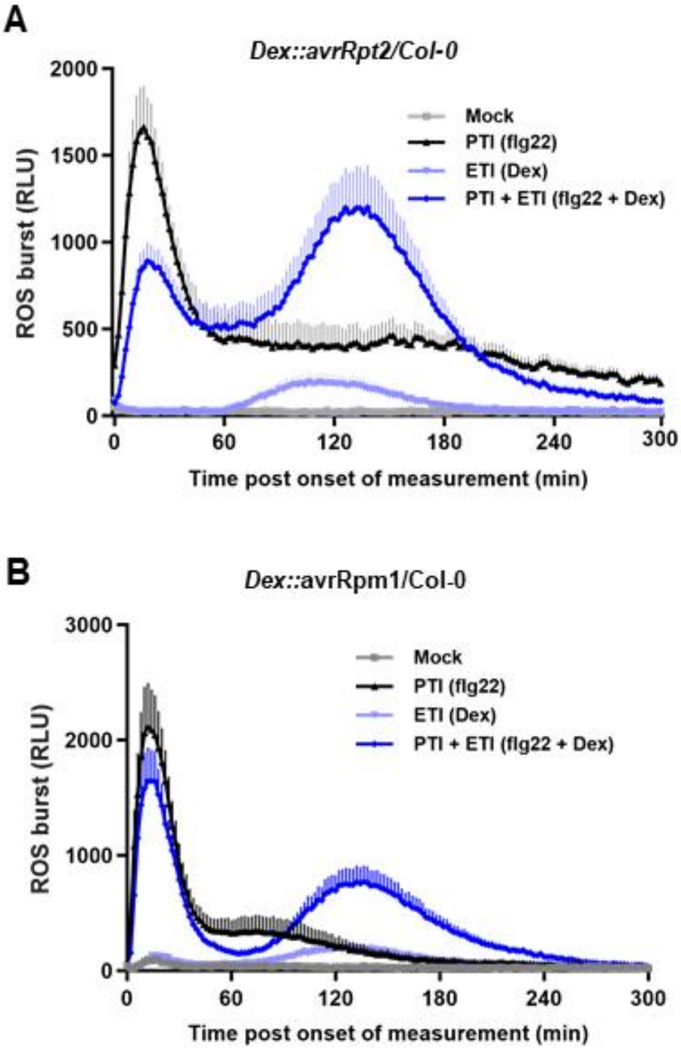


**Figure S5. Valinomycin facilitates more mutual potentiation of PTI and ETI in *Arabidopsis*.**

**(A, B)** Patten-triggered immunity (PTI)- and effector-triggered immunity (ETI)-mediated ROS burst in transgenic plants. (A) *Dex::avrRpt2/*Col-0 and (B) *Dex::avrRpm1/*Col-0 leaves were treated with flg22 and dexamethasone for luminol assay to monitor PTI- and ETI-mediated ROS burst (Mean + SE, n = 12).

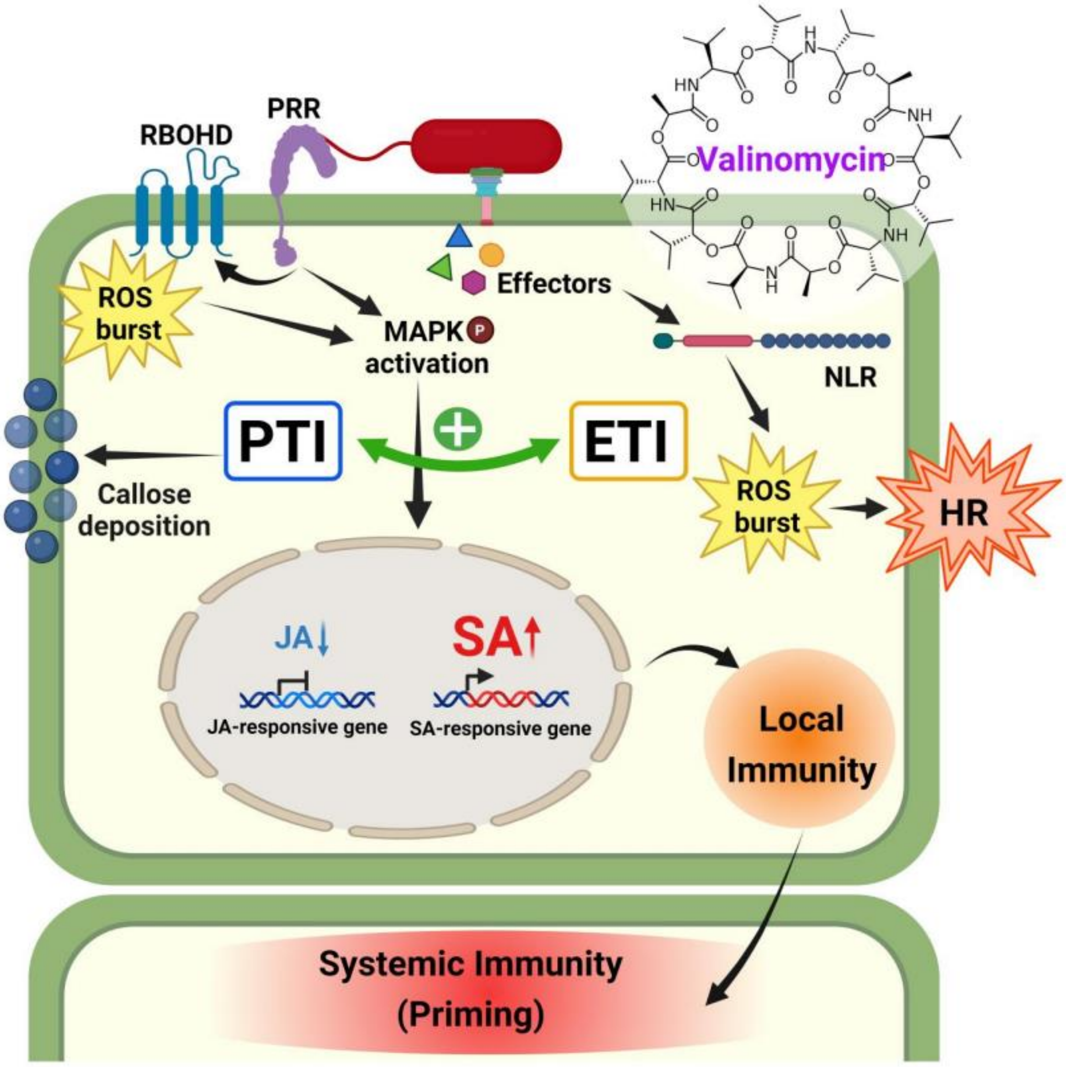


**Figure S6. Working hypothesis of valinomycin-induced local and systemic plant immunity.**

Valinomycin induces robust plant immune responses by potentiating two-tiered innate immune system. Valinomycin strengthened flg22-triggered pattern-triggered immunity (PTI) responses, including ROS burst, MAPK activation, and callose deposition. Second tier of immunity, effector-triggered immunity (ETI), was enhanced by valinomycin pretreatment, which was monitored by quantifying the severity of hypersensitive response (HR). Valinomycin reinforces mutual synergism between two tiers of plant immunity (green arrow). Eventually, local immune-boosting resulted in boosting systemic immunity, thereby valinomycin led to suppression of bacterial growth both locally and systemically. Valinomycin-induced immune-priming is likely to be dependent on jasmonic acid (JA), downregulating JA signaling pathway and then upregulating salicylic acid (SA) pathway. The precise mechanism in terms of hormonal signaling and local/systemic plant immunity needs to be investigated further.
